# Whole-brain meso-vein imaging in living humans using fast 7 T MRI

**DOI:** 10.1101/2025.03.23.644588

**Authors:** O.F. Gulban, R. Stirnberg, D.H.Y. Tse, A. Pizzuti, K. Koiso, M.E. Archila-Melendez, L. Huber, Saskia Bollmann, R. Goebel, K. Kay, D. Ivanov

## Abstract

Non-invasive measurement of the human brain’s angioarchitecture is critical for understanding the basis of functional neuroimaging signals, diagnosing cerebrovascular diseases, and tracking neurodegeneration. Ultra-high-field magnetic resonance imaging (MRI) has achieved mesoscopic (*<* 0.5 mm) imaging of angioarchitecture, revealing fine vascular details that were previously inaccessible in vivo. However, current mesoscopic MRI methods for imaging angioarchitecture face two major limitations. First, acquisition times are prohibitively long, often exceeding 40 minutes, making integration into everyday clinical practice and research projects impractical. Second, even with data successfully acquired, conventional data visualization methods, such as 2D slice browsing and 3D vessel segmentation renders, are rudimentary and have limited effectiveness for navigating and interpreting the complex vascular network. In this paper, we present a fast whole-brain MRI protocol that provides robust images of the brain’s venous network at 0.35 mm resolution in under seven minutes. Additionally, we introduce novel data processing and visualization techniques that enable identification of specific vessel types and more informative navigation of the complex vascular network. We demonstrate that, with these advancements, we can reproduce, in vivo and without intravenous contrast application, the seminal postmortem vasculature images of Duvernoy and Vannson (1999). Furthermore, leveraging the ability for MRI to cover the entire brain, we achieve, for the first time, whole-brain intracortical mesoscopic vein maps in humans. Our acquisition and post-processing methods lay the groundwork for detailed examination of vascular organization across individuals, brain regions, and cortical layers. More generally, these methods make mesoscopic imaging of angioarchitecture viable for broad neuroscientific and clinical applications.

## 1 Introduction

The brain relies more on continuous blood flow than any other tissue, with even brief reductions causing unconsciousness and prolonged deficits leading to irreversible damage (Scharrer, 1960). Therefore, the vascular network (angioarchitecture) is a critical aspect of brain structure, function, and pathology. The vascular network not only delivers oxygen and nutrients but also shapes key physiological and neuroimaging signals (Polimeni, 2025). For example, in functional magnetic resonance imaging (MRI), the blood-oxygenation-level-dependent (BOLD) signal is predominantly influenced by venous blood (Ogawa, Lee, Kay, & Tank, 1990; Ogawa, Lee, Nayak, & Glynn, 1990), which makes the venous angioarchitecture particularly relevant for accurately interpreting the fMRI signal Harel et al. (2010), Kim and Ogawa (2012), Menon (2012), Turner (2002), and Yu et al. (2016). Beyond neuroimaging, venous architecture is important in cerebrovascular health and disease (De Cocker et al., 2018; MacDonald & Frayne, 2015). The organization of small veins is crucial for glymphatic waste clearance (Naganawa et al., 2024), and high-resolution venous imaging aids in detecting small vessel diseases (Van Den Brink et al., 2023; Zwanenburg & Van Osch, 2017) and diagnosing neurodegenerative disorders and cerebral venous thrombosis (Meckel et al., 2010).

Recent advancements in mesoscopic imaging (*<* 0.5 mm) with ultra-high-field MRI have enabled the capture of fine intracortical vascular details, revealing mesoscopic veins within the cortical gray matter (Gulban et al., 2022). This capability stems from the advantages of 7 Tesla MRI, where high spatial resolution and shorter transverse relaxation times of deoxygenated blood enhance sensitivity to venous structures (Ugurbil, 2014). The superior T_2_*-weighted venous contrast at 7 T, compared to 3 T and 1.5 T, has been well established (Koopmans et al., 2008). However, despite these advancements, whole-brain mesoscopic venous imaging in humans remains constrained by long acquisition times, often ranging from 20 to 40 minutes (Federau & Gallichan, 2016; Mattern et al., 2018; Stucht et al., 2015; Van Gelderen et al., 2023), which is impractical due to participant compliance and susceptibility to motion artifacts. In addition to acquisition speed, the analysis and interpretation of complex vascular networks remains a formidable challenge. Despite advancements in mesoscopic vascular imaging (Bernier et al., 2018; Bollmann et al., 2022; Huck et al., 2019; Lü sebrink et al., 2021; Mattern et al., 2018), there has been limited progress in image analysis: current methods are vessel-type agnostic, making it difficult to disentangle vessels and navigate their network structure **Figure 1**).

**Figure 1:**
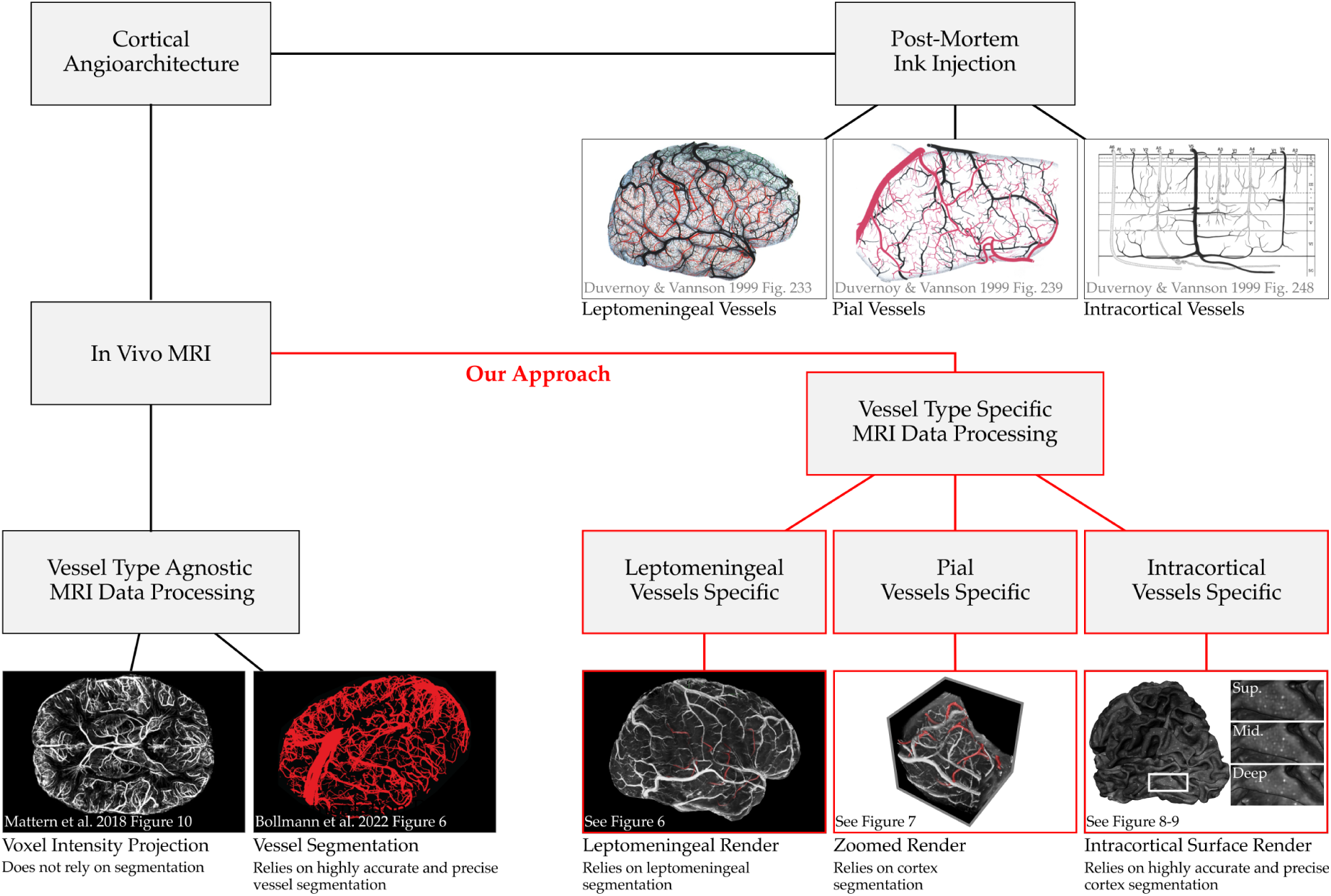
Approach for reconstruction of vasculature images. In this figure, we compare seminal postmortem vasculature images of Duvernoy and Vannson (1999) (upper right), existing vascular imaging methods (lower left), and our approach for reconstructing vascular images (lower right). Based on meticulous post-mortem ink injections and dissections, Duvernoy was able to systematically divide the vasculature network into different vessel types (leptomeningeal, pial, and intracortical vessels). Current MRI-based methods for vascular imaging either provide limited insight (voxel intensity projection) or are overly laborious (vessel segmentation); and in both cases, specific vessel types are not distinguished. Our approach for vasculature imaging relies on simpler and more robust segmentations and exploits modern volume rendering techniques. The resulting visualizations (leptomeningeal render, zoomed render, intracortical surface render) successfully disentangle vessel types and approximate Duvernoy’s vasculature images.

The analysis and interpretation of cerebral vessels is challenging due to their intricate, space-filling architecture. Cerebral vessels traverse the brain’s surface, penetrate deep into the gray matter, and form a multiscale network with diverse spatial connectivity and distribution patterns (Browning, 1884; Duvernoy et al., 1981; Pfeifer, 1940; Salamon, 1971; Scharrer, 1960). A solution to disentangling this complex vascular network is the application of vessel-type-specific data processing and visualization. For example, Duvernoy and Vannson (1999) introduced a framework that categorizes vessels into three distinct types: (1) leptomeningeal, (2) pial, and (3) intracortical vessels (see **Figure 2 B**). Leptomeningeal vessels are the largest, coursing through the subarachnoid space, sometimes floating millimeters above the cortical surface. Pial vessels branch from this network and adhere closely to the cortex. Intracortical vessels include both the main trunks descending into or ascending from the cortical gray matter and the smaller capillary networks interconnecting them.

**Figure 2:**
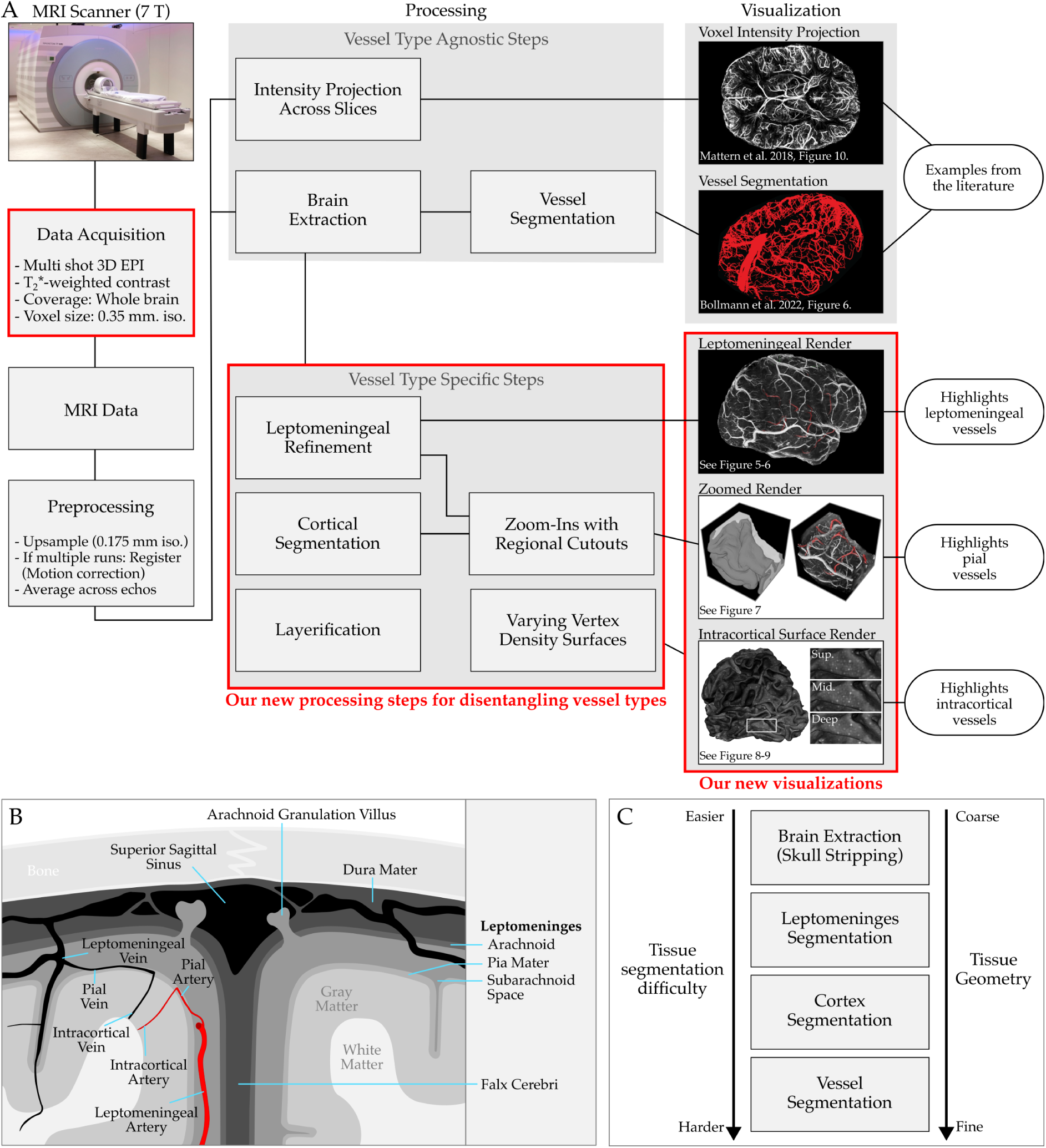
Overview of post-processing. (A) Our data acquisition achieves whole-brain coverage at 0.35 mm isotropic resolution, delivering mesoscopic T_2_*-weighted images in under seven minutes. Our post-processing workflow makes it easy to navigate the vasculature network and distinguish different vessel types down to intracortical mesoscopic details. (B) Schematic of cerebral tissues referenced throughout this manuscript. Note that the capillaries connecting intracortical arteries to veins are not drawn. The illustration depicts a coronal slice from the superior midsection of the human brain. (C) Segmentation difficulty is increased for finer-scale tissue geometry.

In this study, we address two major challenges in mesoscopic in vivo human MRI: the long acquisition times required for whole-brain coverage and the difficulty of extracting vascular information and navigating the complex vascular network. Building upon recent advancements in multi-shot 3D echo planar imaging (EPI) (Stirnberg, Deistung, et al., 2024), we present an optimized venous imaging protocol that achieves whole-brain coverage at 0.35 mm isotropic resolution in under seven minutes. This fast acquisition reduces participant demands and minimizes motion artifacts, while also providing strong venous contrast without any intravenous contrast media. To leverage the fine vascular details captured in our mesoscopic images, we introduce vessel-type-specific data processing and visualization techniques for leptomeningeal, pial, and intracortical veins. Our methods avoid the difficulties of direct vessel segmentation by instead relying on easier-to-achieve tissue segmentations, thereby streamlining the analysis and increasing accuracy of results. We show that we can approximately reconstruct Duvernoy and Vannson (1999)’s seminal postmortem vascular maps in vivo. Moreover, we present the first whole-brain intracortical meso-vein maps in living humans—an accomplishment not yet realized by any other human brain imaging study. Our advances lay the foundation for in-depth investigations of vascular organization across individuals, brain regions, and cortical layers. More broadly, they advance the feasibility of mesoscopic angioarchitecture imaging for diverse applications in neuroscience and clinical research.

## 2 Results

We present an in vivo imaging approach for human venous angioarchitecture that combines whole-brain coverage, 0.35 mm isotropic resolution, and a rapid acquisition time of under seven minutes with strong venous contrast achieved without requiring intravenous contrast media (**Figure 2**). Our fast protocol, in combination with participant preselection, alleviates the head motion artifacts, yielding consistently high-quality images (see **Figure 3-4**). Furthermore, we introduce vessel type-specific data processing and visualization techniques that go beyond conventional intensity projections and segmentation-dependent 3D reconstructions (see **Figure 1-2**). We present in vivo reconstructions of Duvernoy and Vannson (1999)’s postmortem images, capturing the leptomeningeal (**Figure 5-6**), pial (**Figure 7**), and intracortical (**Figure 8-9**) venous network at mesoscale.

**Figure 3:**
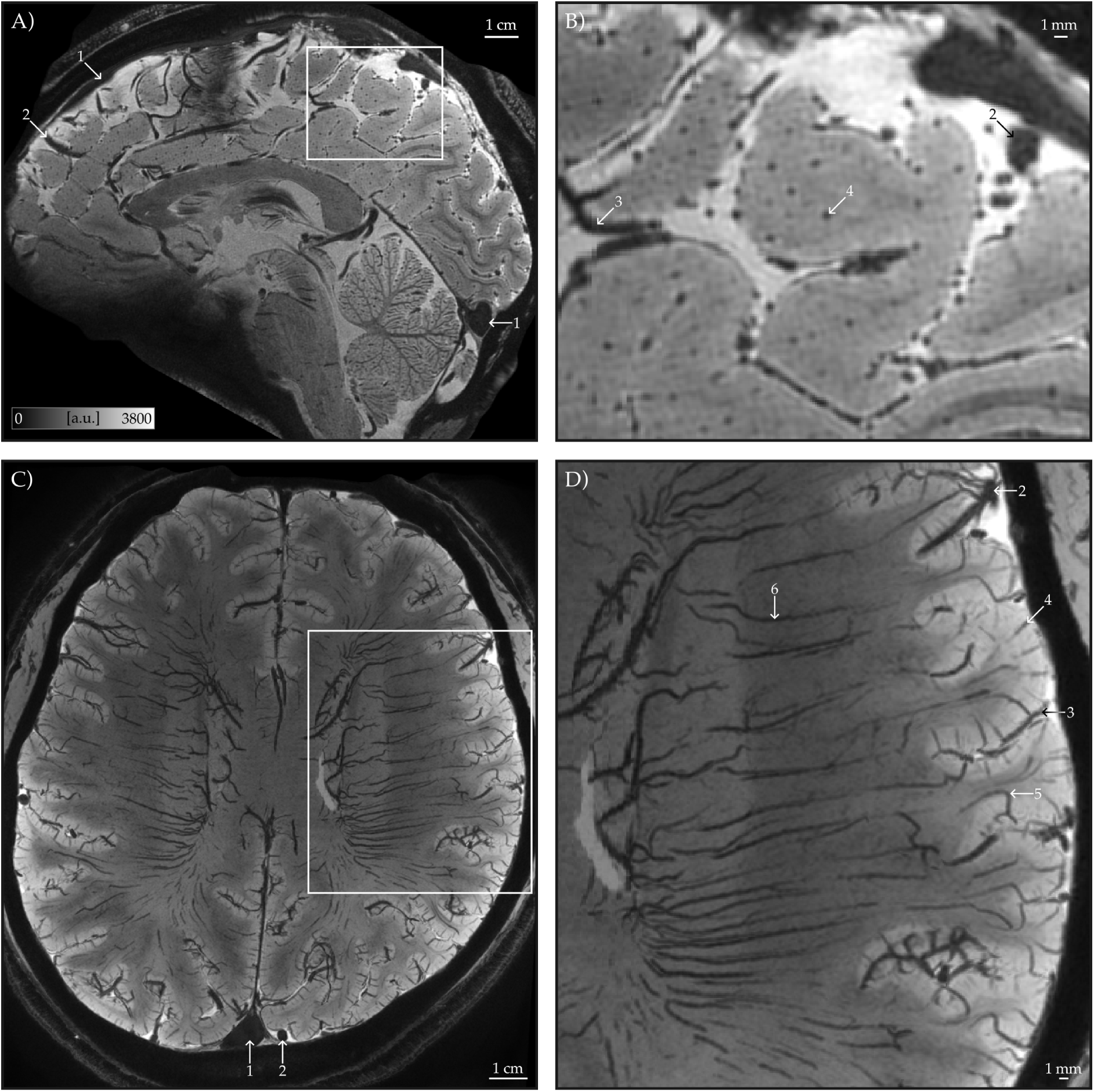
Data quality and venous structures. (A) Our T_2_*-weighted images at 0.35 mm isotropic resolution have high SNR and reveal fine vascular details (shown is the average across all echoes and all four runs of an example participant). To avoid artifacts, we tilted the imaging slab to avoid the eyeballs while still covering the cerebellum and the rest of the cortex. (B) Zoomed-in view highlights intracortical veins. These veins are oriented radially to the cortical surface which is tangentially sectioned in this sagittal slice. (C-D) Minimum intensity projection (3 mm thick) in the transverse plane enhances the visibility of mesoscopic intracortical veins alongside white matter veins. Arrows indicate (1) dural venous sinuses, (2) leptomeningeal veins, (3) pial veins, (4) principal intracortical veins, (5) class V5 intracortical veins, which extend into the white matter, and (6) white matter veins.

**Figure 4:**
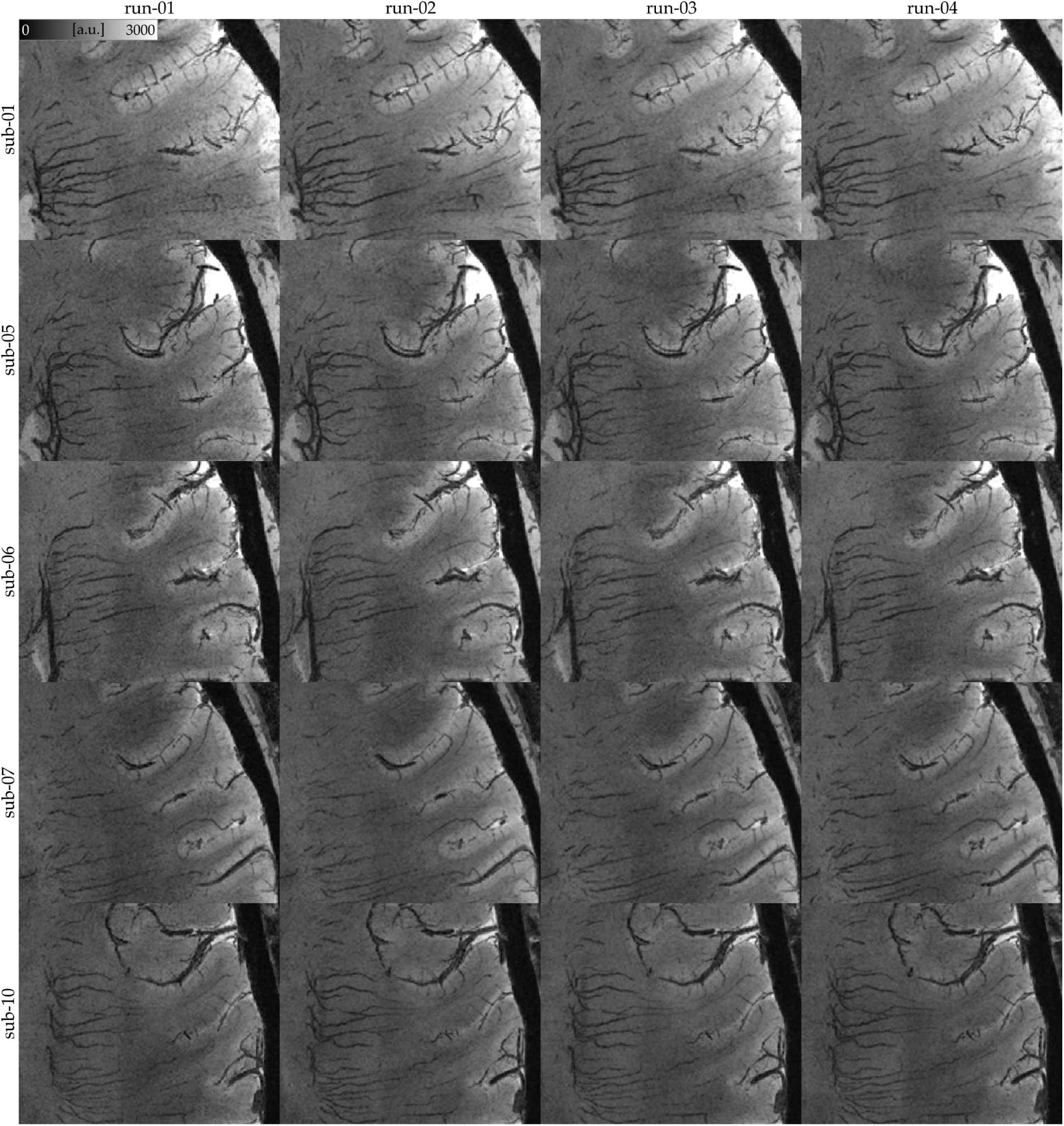
Consistency of echo-averaged T_2_*-weighted images across acquisitions. Each panel presents a minimum intensity projection (3 mm thick) in the transverse plane for a single run. The fine details of mesoscopic veins remain consistently visible across runs, demonstrating reproducibility. All images have been motion-corrected to the first run for direct comparison. The quality of these results suggest that even a single run (under seven minutes) of our multishot multi-echo 3D EPI T_2_*-weighted sequence provides excellent contrast for capturing fine vascular details.

**Figure 5:**
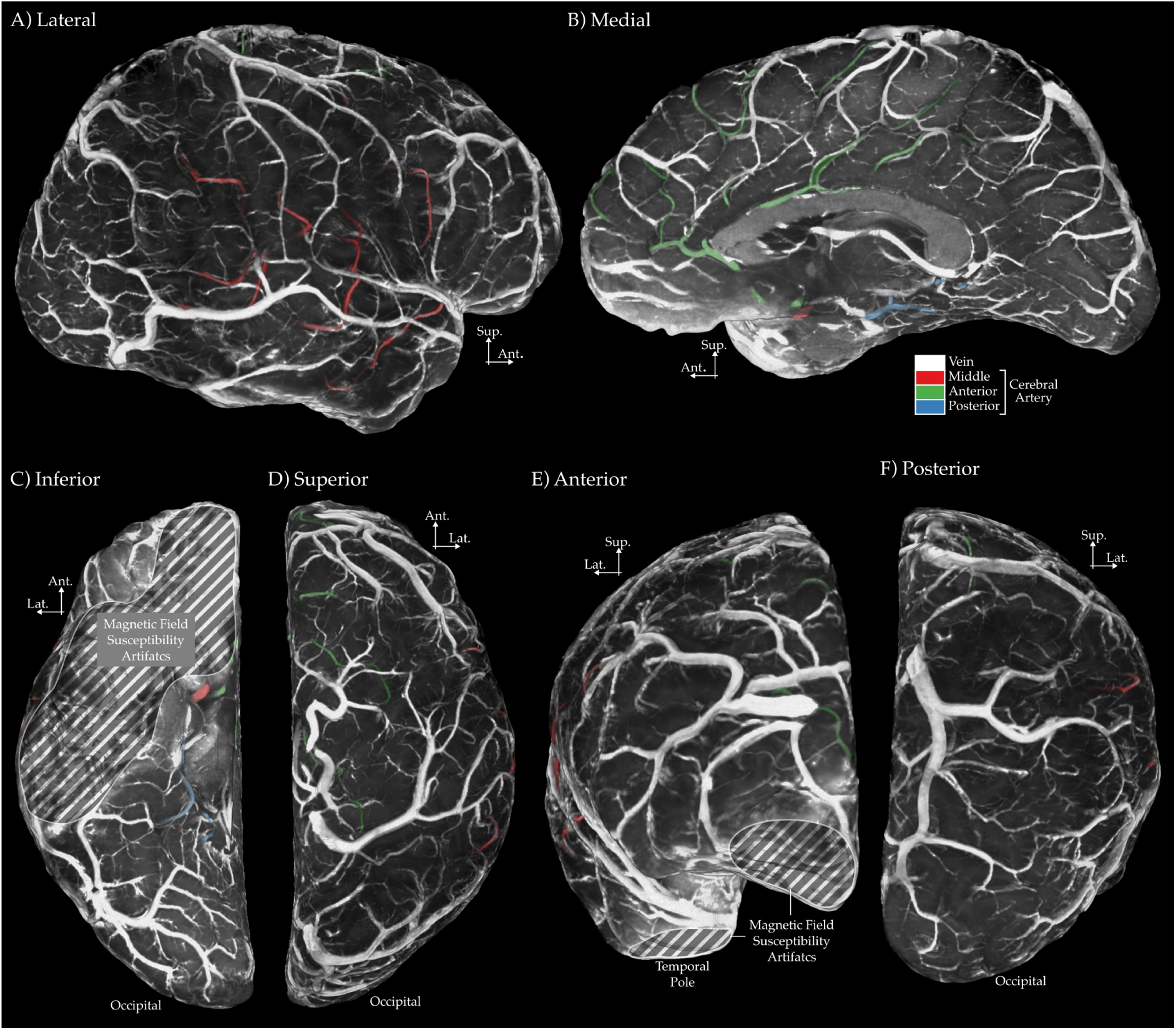
Leptomeningeal angioarchitecture reconstructions using in vivo MRI. Each panel corresponds to an image in Duvernoy and Vannson (1999): (A) lateral view is reconstruction of Figure 233, (B) medial view is a reconstruction of Figure 234, (C) inferior view is a reconstruction of Figure 235, (D) superior view is a reconstruction of Figure 236, (E) anterior view is a reconstruction of Figure 237, (F) posterior view is a reconstruction of Figure 238. Note that the images show “1/T_2_*-weighted” contrast. Smaller veins require a more zoomed in view to be visible on our in vivo images (see Figure 7). In addition, we have marked the inferior brain regions affected by magnetic field susceptibility artifacts, where tissue segmentation becomes unreliable. See Supplementary Figures 3-4 for side by side comparions to Duvernoy and Vannson (1999) with matching visual style.

**Figure 6:**
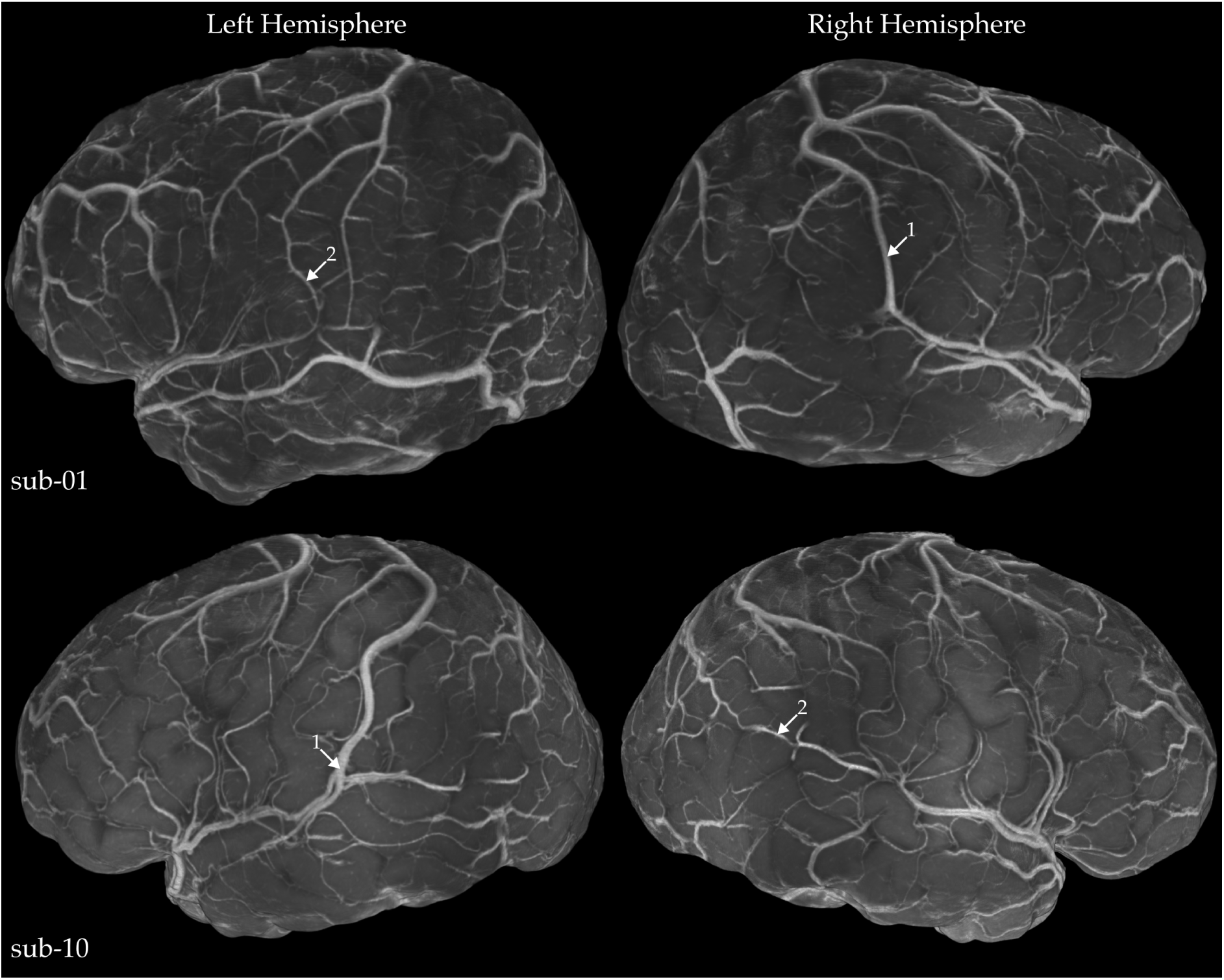
Variation in leptomeningeal veins across hemispheres and individuals. The contrast of these images is ”1/T_2_*-weighted”. Arrows indicate large-diameter anastomotic veins that, while similar at a broad scale, exhibit morphology that is unique to each hemisphere and individual. This highlights the effectiveness of our acquisition and postprocessing methods for detailed anatomical characterization.

**Figure 7:**
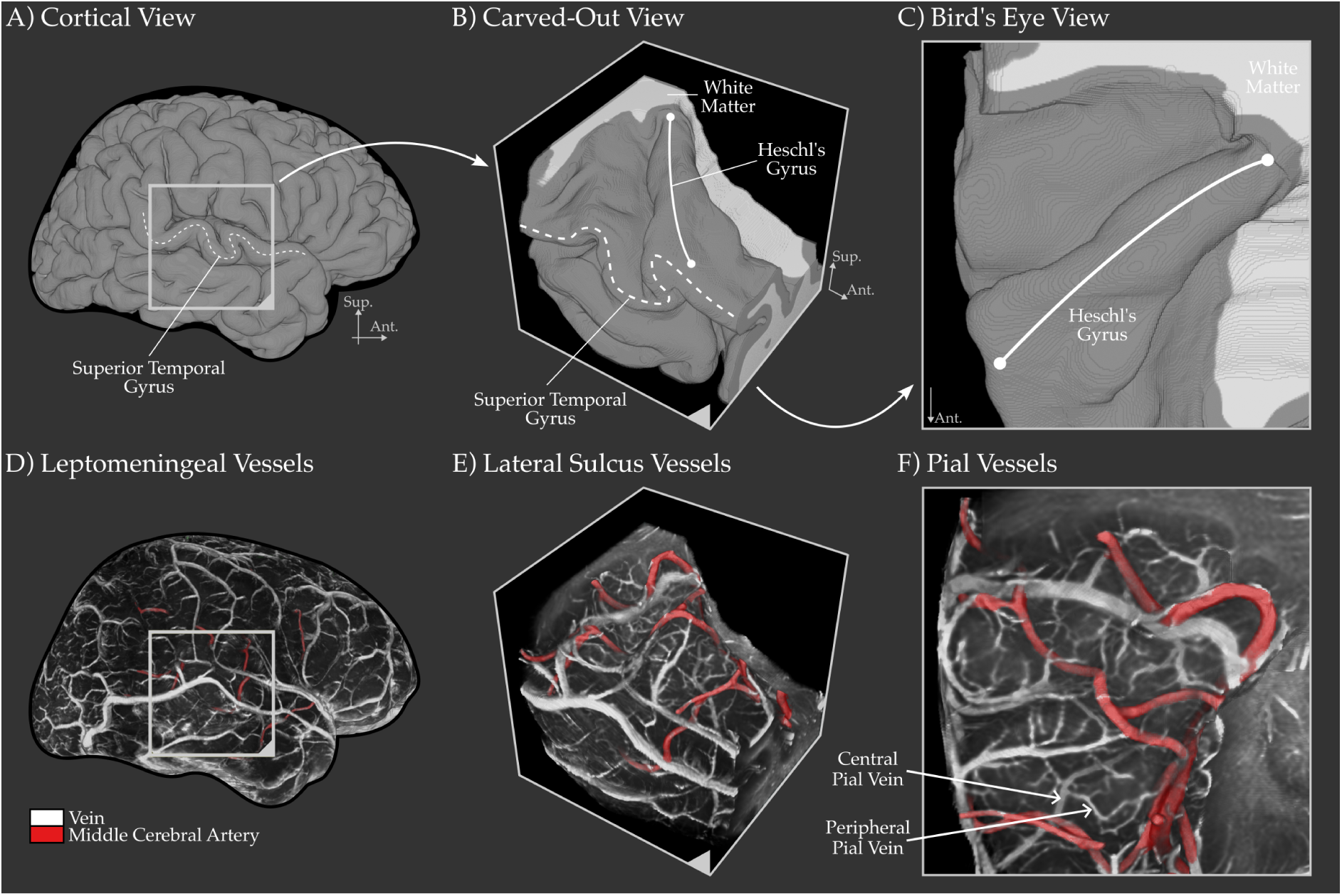
Visualization of pial vessels near the transverse temporal gyri, buried within the lateral sulcus. (A) Segmented gray matter. (B, C) ”Cut-out” view of the temporal lobe after removing the parietal lobe, providing a clearer bird’s-eye perspective of Heschl’s gyrus. (D, E, F) Detailed visualization of the leptomeningeal and pial angioarchitecture within 1.5 mm of temporal cortex gray matter. The middle cerebral artery, colored in red, is classified by tracing its branches towards the brainstem. Examples of central and peripheral pial veins are highlighted with arrows in F. Unlike traditional 2D intensity projections, our 3D visualization enables intuitive navigation and anatomical tracing of the vascular network. This visualization is inspired by Duvernoy and Vannson (1999), Figure 239. See Supplementary Figure 5 for alternative viewing angles.

**Figure 8:**
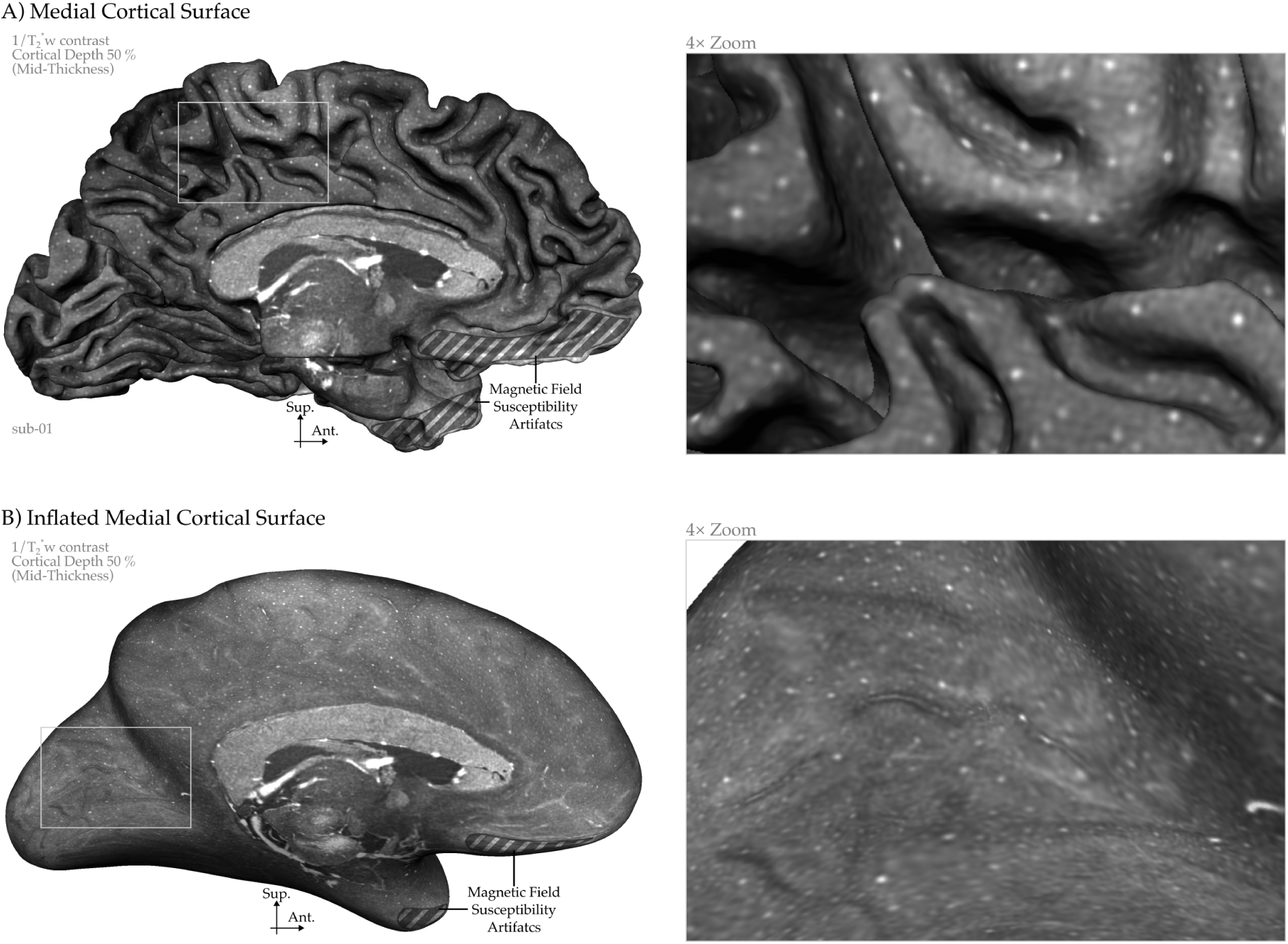
Whole-brain intracortical mesoscopic vein maps. (A) Medial view of middle gray cortical surface (mid-thickness) displaying “1/T_2_*-weighted” contrast. (B) Inflated version revealing vascular patterns within sulci. Mesoscopic intracortical veins appear as bright dots. Inferior brain regions affected by magnetic field susceptibility artifacts, where tissue segmentation becomes unreliable, are indicated.

**Figure 9:**
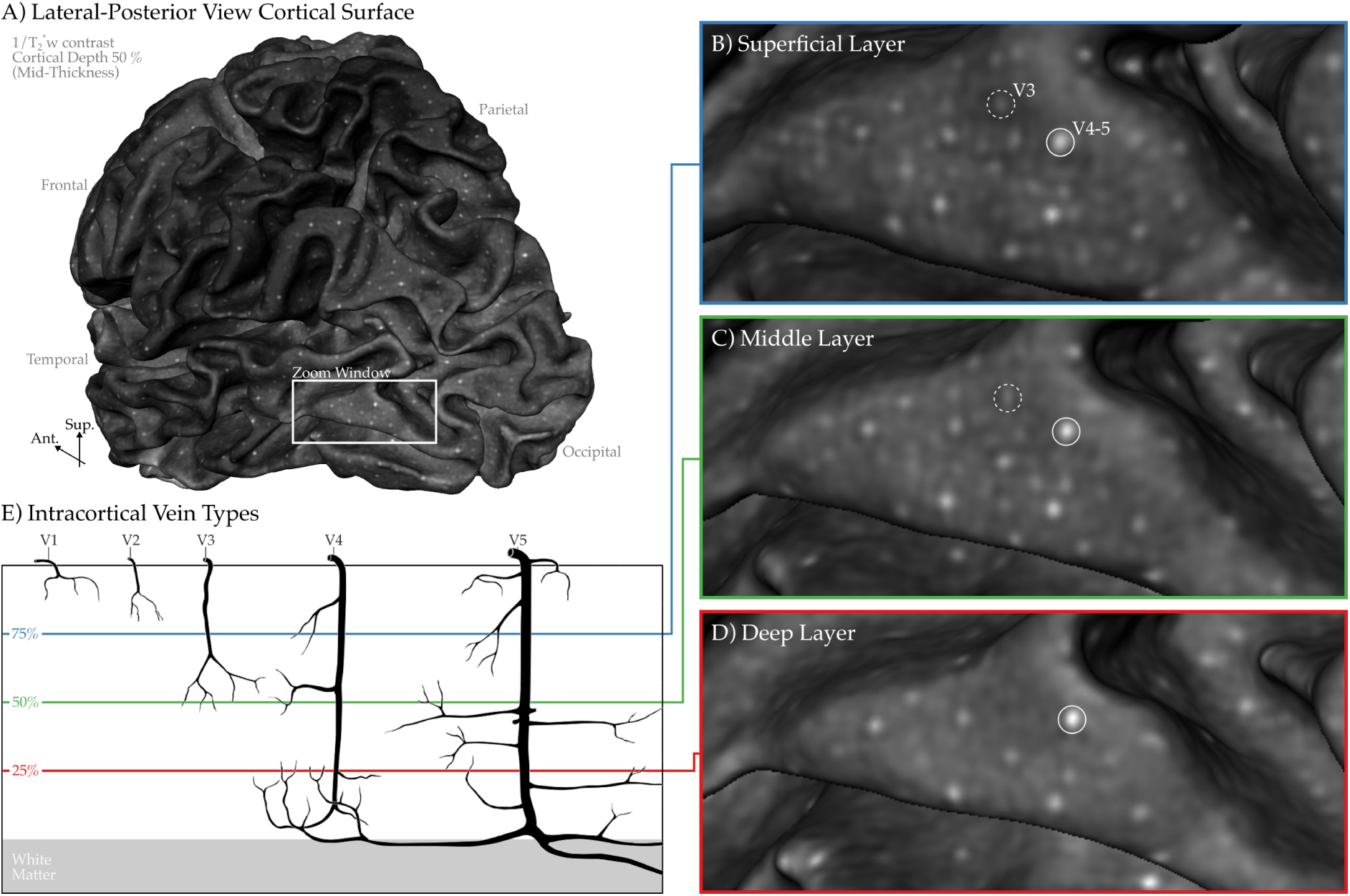
Intracortical veins visualized across different cortical depths in vivo. (A) Middle gray matter surface (mid-thickness) shown from a lateral, angled posterior perspective. (B–D) Geometric cortical layers at 25% (deep), 50% (middle), and 75% (superficial) of the local cortical thickness within the zoomed-in region from Panel A. (E) Schematic is redrawn from Duvernoy and Vannson (1999) Figure 248, and rearranged to isolate intracortical veins. Tracking the appearance and disappearance of bright dots across cortical depths allows us to identify class 3, 4, and 5 intracortical veins (see labeled circles).

### 2.1 Data Quality and Consistency

Our T_2_*-weighted images excel at capturing mesoscopic veins (*<* 0.5 mm in diameter), because of the strong contrast between deoxygenated blood and surrounding tissues at 7 T (see **Figure 3**). Notably, even a single acquisition appears sufficient to distinguish meso-veins from brain tissue (see **Figure 4** and **Supplementary** Figure 1). The high consistency of venous structures across runs and participants suggests that, when head motion is minimized, the available SNR and venous contrast are adequate for reliably identifying meso-veins.

### 2.2 Leptomeningeal Veins and Arteries

Figure 5 compares the vascular details captured in our in vivo MRI data with the anatomical illustrations published in Duvernoy and Vannson (1999). The images clearly demonstrate that our mesoscopic imaging at 0.35 mm isotropic resolution, combined with our leptomeningeal vessel-specific processing and visualization techniques allow us to reproduce the venous vascular network visible in Duvernoy and Vannson (1999)’s postmortem vascular images. We can clearly identify all three major classes: superior, middle, and inferior cortical veins, as well as the anastomotic cortical veins on the lateral surface of the hemispheres. Note that although the smaller pial veins are less apparent than the leptomeningeal veins compared to Duvernoy and Vannson (1999)’s drawings, we address this issue in the ”Pial Veins” subsection.

Additionally, our images enable the identification of large leptomeningeal arteries as well. Interestingly, the dark appearance of the arteries—sometimes even darker than that of the veins—may seem paradoxical, as arterial blood has higher oxygenation levels, leading to slower transverse decay and therefore brighter T_2_*-weighted signal. However, the darkness of the large leptomeningeal arteries is not due to transverse relaxation but rather to the blood motion artifact (Gulban et al., 2022). Arterial blood is known to flow at high speeds within the large arteries, and this motion introduces artifacts due to the time delay between phase encoding and frequency encoding during MRI data acquisition. This artifact manifests as a displacement of the arterial bright signal along the frequency encoding axis, proportional to the velocity of blood flow. A clear visual example of this artifact is presented in Gulban et al. (2022) Supplementary Figure 1. However, because our 3D EPI acquisition scheme employs different spatial encoding and collects MR signal over a longer acquisition window compared to the smaller windows used in, for example in Gulban et al. (2022), the reconstructed image exhibits a different artifact appearance. Instead of a bright spot being displaced near large arteries and leaving the artery itself as a dark structure, the arteries in our images simply appear as dark tubular structures without a surrounding bright artifact. This characteristic dark appearance of large leptomeningeal arteries is particularly important, as veins also appear dark in our T_2_*-weighted images, making it difficult to distinguish arteries from veins based solely on signal intensity. We confirmed that these vessels are arteries not only based on image contrast but also by tracing their trajectories in relation to known anatomical landmarks. Specifically, we followed their paths toward the brainstem, where their continuity with major arterial structures, such as the basilar and vertebral arteries, unambiguously identifies them as arteries. Additionally, we verified our arterial trajectory tracing using MP2RAGE UNI images, in which large leptomeningeal arteries appear as bright voxels, while veins remain invisible.

Figure 6 further highlights variations across hemispheres, brain areas, and individuals in the large superficial anastomosing veins (Tomasi et al., 2022). For instance, in sub-01, the right hemisphere features a large superior anastomotic vein, the vein of Trolard. In contrast, the left hemisphere exhibits two smaller superior anastomotic veins and a dominant inferior vein, the vein of Labbé. Overall, Figure 5**-6** underscore the advantages of our real-time volume rendering in effectively capturing variations in leptomeningeal veins and demonstrate how closely our in vivo T_2_*-weighted mesoscopic MRI data reproduces the original post-mortem hand-drawn illustrations reported in Duvernoy.

### 2.3 Pial Veins

While the whole-brain visualizations in Figures 5-6 highlight the leptomeningeal angioarchitecture, the level of detail in the pial angioarchitecture does not seem to match that of the anatomical illustrations in Duvernoy and Vannson (1999). This discrepancy arises from two main factors: (1) Despite achieving mesoscopic imaging at 0.35 mm isotropic resolution, which is high by today’s standards, our voxel size remains insufficient to capture the thinner branches of the pial vessels. (2) Our current voxel value rendering approach is not optimized to fully reveal the fine details of the pial vasculature.

While we leave the first issue for future imaging optimization efforts (Stirnberg, Gulban, et al., 2024; Stirnberg et al., 2025), we address the second by introducing a refined voxel value rendering technique in Figure 7. To demonstrate its effectiveness, we focus on the superior temporal gyrus (Heschl’s Gyrus Heynckes et al. (2022)), a region that features both large leptomeningeal veins and arteries, along with intricate pial vasculature embedded within a deep sulcus (Gulban (2020) Figure 5.4-5.5). This area is typically invisible from the lateral surface without virtual dissection techniques such as “opening” or “carving out” the sulcus. The refined voxel value rendering reveals distinct large trunks of leptomeningeal arteries and veins. Additionally, it also clearly depicts pial veins. Duvernoy defines two types of pial veins: central and peripheral. We can see both pial vein types in our zoomed cutout voxel value renders (Figure 7). The central pial veins that drain from multiple smaller pial veins toward the lateral surface are visible. The smaller peripheral pial veins are also visible. The peripheral veins can even be traced to their tributaries (the ascending veins). The peripheral veins spread across the pial surface and eventually merge with the central veins, which connect to the larger lateral leptomeningeal veins. Some of these pial veins also seem to exhibit anastomotic connections, further detailing the complexity of the vascular network.

### 2.4 Intracortical Veins

Intracortical veins exhibit both laminar and radial features. The laminar aspect is associated with the density of the capillary network at the microscale, which remains beyond our imaging resolution. However, mesoscopic intracortical veins are undoubtedly visible in our images (Figure 3**-4**). To further support this, Figure 8 demonstrates the cortical mid-thickness surface, revealing veins as distinct dot-like structures distributed across the cortical surface that are oriented radially with regard to the inner and outer boundary of the gray matter. The meso-veins appear as dot-like structures because the thin vessels, with highly curved and branching geometries, often display fractal patterns. When viewed using 2D slice browsing, it’s rare to see the veins as continuous tubes, as the slice is more likely to cut across the vessels at an angle rather than parallel to their trunks.

To our knowledge, this is the first whole-brain visualization of intracortical meso-veins in humans. While intracortical meso-veins can be observed in several previous studies showing 2D brain slices (Budde et al., 2011; Duyn et al., 2007; Kemper et al., 2018; Koopmans et al., 2008; Mattern et al., 2018; Petridou et al., 2010; Sánchez-Panchuelo et al., 2012; Stirnberg, Deistung, et al., 2024; Trampel et al., 2017; Zwanenburg et al., 2011), these 2D slices are insufficient for capturing the full extent of meso-veins across the entire cortical surface. approach requires cortex-specific visualization through surface inflation and/or flattening. Our previous work highlighted these meso-veins in small patches of virtually flattened human cortices (Gulban et al. (2022), Figure 4), and more recently, whole-brain visualization of intracortical meso-veins has been achieved only in primates (Autio et al. (2024), Figure 2).

To further elucidate the intracortical meso-veins, we demonstrate their cortical depth-spanning properties Figure 9. Our variable-density surface reconstructions enable high-fidelity visualization of these veins and tracing across different cortical depths. As seen in the figure, relatively large Class 4 and 5 veins (V4, V5) are easily identifiable, while some veins vanish in the mid-thickness of the cortex, potentially corresponding to Class 3 veins (V3). While Duvernoy provides specific size measurements for these veins, these values may be unreliable, as he notes that post-mortem methods do not account for the actual in vivo diameters of veins filled with blood (see Duvernoy et al. (1981), and Gulban et al. (2022), Supplementary Figure 9). We also refrain from interpreting the diameters of these meso-veins due to a different artifact present in our T_2_*-weighted images. The blooming artifact, caused by deoxygenated blood and the orientation of the vein relative to the main magnetic field (B0), introduces magnetic field susceptibility distortions, making veins appear larger than their true size (Ogawa, Lee, Kay, & Tank, 1990). The dark appearance of intracortical veins is particularly pronounced at 7 T due to the faster transverse decay of deoxygenated blood (Koopmans et al., 2008). While this phenomenon is an artifact of our imaging method, we think that it helps us to identify the meso-veins more easily using voxels larger than their actual trunks. Given the high curvature of the human cortex, it is unlikely that any vein aligns perfectly with the main magnetic field, therefore it is reasonable to assume that most veins experience the blooming effect, further darkening the surrounding tissues.

## 3 Discussion

In this study, we have demonstrated data acquisition and visualization methods for studying cortical venous angioarchitecture using whole-brain mesoscopic T_2_*-weighted MRI at 7 T. While previous attempts have been made to acquire whole-brain data at similar resolutions (Federau & Gallichan, 2016; Mattern et al., 2018), we have achieved a substantial speed-up, reducing acquisition times to under seven minutes using multi-shot 3D EPI (Stirnberg, Deistung, et al., 2024). However, efficiently acquiring high-quality data is only part of the challenge. Many MRI-based vascular mapping studies remain constrained by slice-wise intensity projections or 3D model reconstructions that are heavily dependent on vessel segmentation quality. In contrast, we introduce a novel approach for visualizing the human venous angioarchitecture (Figure 2).

### 3.1 Considerations for fast mesoscopic vein imaging

Using multi-shot, multi-echo 3D EPI, we achieved whole-brain coverage at 0.35 mm isotropic resolution in just under seven minutes per scan. This acquisition time falls well within the range of routine clinical protocols, reducing the immediate need for additional motion correction strategies. While prospective motion correction techniques hold great promise for minimizing head motion (Federau & Gallichan, 2016; Maclaren et al., 2012; Stucht et al., 2015; Van Gelderen et al., 2023), they have yet to be fully integrated into daily clinical practice due to technical and logistical challenges. Our results demonstrate that high-quality mesoscopic vascular imaging can already be achieved within clinically feasible scan times without relying on these additional tools (see Figure 4 **and Supplementary** Figure 1). Looking ahead, further refinements to acquisition protocols may shorten scan durations even more, complementing future developments in motion correction and broadening the accessibility of mesoscopic angioarchitecture imaging for both research and clinical applications.

### 3.2 Vessel type specific post-processing

We employed two distinct 3D data visualization frameworks to depict the cerebral blood vessels captured in our images. For the leptomeningeal and pial vessels, we utilize real-time voxel value rendering in combination with the cortical vasculature dissection strategy used by Duvernoy et al. (1981). This synthesis of voxel value rendering with traditional dissection techniques was chosen because it effectively reconstructed the widely influential hand-drawn illustrations by Jean-Louis Vannson (Duvernoy et al., 1981; Duvernoy & Vannson, 1999). These plots highlight the complex space-filling properties of vessels suspended within the subarachnoid space, adjacent to or near the pia mater, providing an optimal visualization of the leptomeningeal and pial vasculature.

While brain imaging techniques have advanced significantly since the 1980s and 1990s—when Duvernoy and Vannson produced their seminal vascular maps—visualization methods for angioarchitecture have lagged behind, often failing to match the clarity and precision of their work. The most common approaches today rely on ”vessel type agnostic” 2D voxel intensity projections or 3D voxel label renders (Bernier et al., 2018; Bollmann et al., 2022; Huck et al., 2019; Koopmans et al., 2008; Lü sebrink et al., 2021; Mattern et al., 2018; Stirnberg, Deistung, et al., 2024). While both methods serve important purposes, they also have limitations. 2D voxel intensity projections lose depth and directional information, with greater information loss as projection window size increases. 3D voxel label renders can preserve depth and directional cues through lighting and surface reflectance, but are highly dependent on the accuracy of vessel segmentation. Achieving accurate vessel segmentation is extremely challenging due to the fine and complex geometry of vessels. Manual or semi-automated segmentation of high-resolution vascular datasets is highly time-consuming (Bollmann et al., 2022), and the development of more efficient automated vascular segmentation algorithms is still an active research area (Xu et al., 2024). To address these challenges, our vessel type-specific processing and visualization techniques provide a practical middle ground (see Figure 2). Instead of relying on demanding vessel segmentations, users perform simpler, more reliable tissue segmentations and leverage voxel value rendering to directly visualize vascular structures. For instance, widely used brain extraction methods can be followed by leptomeningeal refinement to enable visualization of leptomeningeal vessels. Currently, this refinement is performed manually, but it presents an opportunity for further automation, potentially using artificial intelligence methods.

For intracortical vessels, we employ triangular mesh reconstructions with one-to-one voxel-to-vertex sampling. Crucially, instead of the conventional approach of expanding a reference mesh along vertex normals by a fixed percentage of local cortical thickness, we use varying vertex density meshes. This modification enables a more accurate representation of fine intra-cortical details across different cortical geometric layers (Gulban & Huber, 2024) and mitigates aliasing artifacts during the projection and interpolation of MRI data onto surface meshes (Polimeni et al., 2018). The effectiveness of our approach is evident in Figure 9, where single-voxel-wide veins are clearly identifiable across cortical depths. While our current work focuses on intracortical veins visualized with T_2_*-weighted imaging, our varying vertex density meshes can be readily applied to other meso-vessel imaging modalities, such as time-of-flight (TOF) angiography (Bollmann et al., 2022; Lü sebrink et al., 2021; Stucht et al., 2015) or previously acquired mesoscopic datasets (Federau & Gallichan, 2016; Lü sebrink et al., 2021; Mattern et al., 2018). Additionally, although our intracortical meso-vein visualizations were initially inspired by Autio et al. (2024) (Figure 2), our vessel-type-specific advancements may, in turn, enhance non-human imaging (e.g., (Autio et al., 2024; Bolan et al., 2006)), facilitating clearer reconstructions of pial and leptomeningeal vasculature, and extraction of information across cortical layers.

### 3.3 Conclusions and Future Directions

Our work marks a significant step toward practical, high-fidelity mesoscopic venous imaging in vivo, yet exciting opportunities for further advancements remain. Future improvements in imaging protocols and MRI hardware could enhance both spatial resolution and venous contrast while maintaining clinically feasible scan times (Stirnberg, Gulban, et al., 2024) or accelerate the proposed imaging protocol even further (Stirnberg et al., 2025). Additionally, while our vessel-type-specific processing and visualization techniques allow navigation of the vascular network without requiring vessel segmentation, access to accurate vessel segmentation would enable deeper quantitative analyses of vascular networks, including connectivity, tortuosity, and regional perfusion patterns (Xu et al., 2024). Beyond venous imaging, integrating mesoscopic arterial imaging could provide a more comprehensive view of cortical angioarchitecture (Bollmann et al., 2022), bringing in vivo vascular reconstructions closer to the seminal work of Duvernoy and Vannson (1999). Although future MRI hardware and sequence advancements may surpass the imaging methods presented here, our vessel-type-specific processing and visualization techniques will remain valuable for studying mesoscopic cortical angioarchitecture. By further streamlining acquisition, processing, and analysis, mesoscopic neurovascular imaging will continue to evolve as a powerful tool for investigating cerebrovascular health, neurovascular coupling, and pathophysiology in living humans.

## 4 Methods

### 4.1 Participants

Five healthy participants (1 female, aged 25–38 years) were recruited for this study, consisting of a one-hour MRI session at 7 Tesla. Participants were selected based on their prior experience with 7 Tesla MRI experiments and their ability to remain still for extended periods, which is an essential factor in minimizing the bulk head motion artifacts for high-resolution imaging. Similar preselection strategies have been employed in previous high-resolution MRI studies (Allen et al., 2022; Bollmann et al., 2022; Gulban et al., 2022; Tardif et al., 2015). Informed consent was obtained from all participants prior to the experiment. The study was approved by the research ethics committee of the Faculty of Psychology and Neuroscience of Maastricht University and experimental procedures followed the principles expressed in the Declaration of Helsinki.

### 4.2 Data Acquisition

We acquired T_2_*/susceptibility-weighted whole-brain images using a multi-shot, multi-echo 3D echo planar imaging (3D EPI) sequence on a 7 T MRI scanner (Siemens Healthineers, Magnetom “7TPlus”) equipped with a 32-channel receive-only head coil (Nova Medical). Specifically, we employed a modified Skipped-CAIPI 3D-EPI sequence (Stirnberg & Stö cker, 2021), which optimizes both scan time and signal-to-noise ratio (SNR) by using a longer repetition time (TR) than conventional approaches (Stirnberg, Deistung, et al., 2024). At 7 T, venous contrast remains superior to 3 T for echo times up to 40 ms (Koopmans et al., 2008), yet current 7 T T_2_*-weighted venous imaging has been limited to echo times of 18 ms (Federau and Gallichan (2016), Huck et al. (2019), Mattern et al. (2018), Stirnberg, Deistung, et al. (2024), and Stucht et al. (2015), however Van Gelderen et al. (2023) goes up to 39 ms). To maximize venous contrast, we extended our echo times beyond this threshold and incorporated multiple echoes. Compared to the 3D gradient-recalled echo (3D GRE) technique used in our previous mesoscopic imaging study (Gulban et al., 2022), multi-shot multi-echo 3D EPI provides four times larger coverage, from a slab to the whole brain, while maintaining the same high spatial resolution at 0.35 mm isotropic voxel, in less than 7 minutes scanning time. For B0 shimming, we have used an in-house developed workflow (Tse et al., 2016). In addition, we have also acquired a 0.7 mm isotropic resolution MP2RAGE image (Marques et al., 2010) for T1 weighted reference. Our 3D EPI parameters were as follows: nominal voxel resolution = 0.35 × 0.35 × 0.35 mm³, TR = 52.5 ms, TE1–3 = [9.46, 24.66, 39.86] ms, flip angle (FA) = 10°, matrix size = 572 × 572 × 380 voxels, field of view (FOV) = 20 × 20 × 13.3 cm³, parallel imaging acceleration = 3 × 2, segmentation/EPI factor = 40/5, and total volume acquisition time = 6 min 48 sec. **Supplementary** Figure 1 illustrates the positioning of the imaging slab, which was tilted to approximately align with the anterior commissure–posterior commissure (AC–PC) plane. This positioning allowed us to include the entire cerebellum within the imaging volume while also strategically shifting the “top of the head” bone and fat-wrapping artifact toward the nasal cavity, where its impact on overall image quality is significantly reduced. Four consecutive 3D EPI runs were acquired. The frequency encoding polarity was flipped in every other scan to eliminate minor segmentation artifacts near most severely off-resonant brain areas by final averaging of the magnitude images following between-run head motion correction (Stirnberg, Deistung, et al., 2024). Importantly, we found that allowing participants a brief break between runs—explicitly instructing them, ”Please feel free to slightly move your head if needed”—significantly reduced bulk head motion during subsequent acquisitions. In our experience, this approach yielded higher image quality compared to the conventional instruction to ”stay as still as possible throughout the entire scanning session”.

Our MP2RAGE parameters were as follows: nominal voxel resolution = 0.7 × 0.7 × 0.7 mm3, TR = 5000 ms, TE = 2.47 ms, TI1-2 = [900, 2750] ms, FA1-2 = [5°, 3°], 320 × 320 × 240 voxels, 22.4 × 22.4 × 16.8 cm3 slab dimensions, GRAPPA = 3, Partial Fourier = 6/8, Bandwidth = 250 Hz/Px, and 8 minutes duration.

Further details for both 3D EPI and MP2RAGE acquisitions can be found in our protocol documents (https://doi.org/10.5281/zenodo.14145584).

### 4.3 Data Analysis

#### 4.3.1 Bulk head motion correction

There are two primary types of bulk head motion to consider: (i) within-run head motion, which introduces blur and ringing artifacts, and (ii) between-run head motion, which affects image averaging and reduces overall data quality . To mitigate within-run head motion, we preselected participants with prior experience in high-field MRI studies and familiarity with minimizing head movement. This participant preselection strategy has been successfully implemented in previous studies (Allen et al., 2022; Bollmann et al., 2022; Gulban et al., 2022; Tardif et al., 2015). Additionally, we kept individual run durations to under seven minutes, as scan durations exceeding 20 minutes typically require prospective motion correction (Lü sebrink et al., 2017; Mattern et al., 2018; Stucht et al., 2015) or fat-based motion navigators (Federau & Gallichan, 2016; Gallichan et al., 2016; Van Gelderen et al., 2023). Our shorter acquisitions, combined with participant preselection resulted in excellent data quality where we did not discard any data upon quality assessment (see Figure 3**-4**).

To correct for between-run motion, we implemented a multi-step preprocessing pipeline:

- **Cropping:** Non-cortical areas of the images were cropped to reduce computational demands, particularly RAM usage, in subsequent processing steps.
- **Upsampling:** Individual echos were resampled to 0.175 mm isotropic resolution using cubic interpolation for better preserving fine structural details in the upcoming steps (Allen et al., 2022; Wang et al., 2022).
- **Echo averaging:** Signal intensities were averaged across all echoes for each voxel, improving signal-to-noise ratio (SNR) and generating a reference image for motion estimation.
- **Motion correction:** A rigid-body transformation (6 degrees of freedom) was applied to align all runs to the first run, using the reference image. Each individual echo image was then corrected via linear interpolation using ITK-SNAP v4.2.2 and c3d (Yushkevich et al., 2006).
- **Final averaging:** After motion correction, all runs and echoes were averaged, yielding a final brain image with a nominal resolution of 0.175 mm isotropic voxels.

Note that, given that even single runs provide high-SNR data (see Figure 4 **and Supplementary** Figure 1), steps for the motion correction can be omitted. In such cases, we recommend upsampling and averaging across echoes to maintain data quality.

#### 4.3.2 Tissue Segmentation

Our goal was to achieve high-precision tissue segmentation optimized for comprehensive visualization of leptomeningeal vessels, pial vessels, and intracortical vessels across the whole brain. To accomplish this, we first processed the MP2RAGE UNI images using the BrainVoyager v24.0 (Goebel, 2012) advanced segmentation pipeline, following default parameters and procedures to generate initial segmentations for the brain mask, gray matter mask, and white matter mask. Next, we registered the 0.7 mm isotropic MP2RAGE data to our preprocessed 0.175 mm isotropic EPI data using semi-automatic non-linear registration (using ITSNAP and greedy (Yushkevich et al., 2006)). Once the registration parameters were estimated, we resliced the initial segmentation masks to 0.175 mm isotropic EPI space using an interpolation method specifically designed for reslicing and registering tissue label files. After transforming the segmentation masks into EPI space, we performed manual edits to improve the precision and accuracy of each tissue type in ITKSNAP. Specifically, we filled in the subarachnoid space, avoiding the dura mater, arachnoid granulation villi, and subdural venous sinuses (see Figure 2 **B**). We have ensured smoothness of the segmented tissues by applying the LN2 RIM POLISH program from LayNii upon the completion of manual edits. The result of the leptomeningeal refinement segmentation is visible in **Supplementary** Figure 2. We emphasize that achieving high-accuracy and high-precision segmentation for all three target tissues -leptomeninges, cortical gray matter, and cortical white matteris critical for the reliability of our subsequent analyses and visualizations.

### 4.4 Data visualization

#### 4.4.1 Voxel value rendering for leptomeningeal and pial vessels

We begin by applying a reciprocal transformation to our T_2_*-weighted images, where each voxel intensity is transformed as “1 / T_2_*-w”, with division-by-zero cases set to zero. This simple yet effective transformation inverts vessel contrast, making cerebral blood vessels appear bright instead of dark, thereby enhancing their natural appearance while simultaneously dimming the bright cerebrospinal fluid (CSF) and gray matter. After this transformation, we convert the image precision from float32 to unsigned integer 8 (uint8) by clipping intensity values at the 1st and 99th percentiles to enhance contrast and preserve dynamic range. For visualization, we utilize ”real-time volume rendering” in BrainVoyager v24.0 Goebel (2012), applying the following parameters: step size = 1.0, absorption = 1.0, LUT range = [0, 225], cubic sampling = on, keep large gradients along rays = off, use light for shading = off.

#### 4.4.2 Varying vertex density triangular meshes for intracortical vessels

Visualization of intracortical vessels is performed in two main steps. First, we compute equidistant geometric layers using the LayNii v2.7.0 (Huber et al., 2021) LN2 LAYERS program with the parameter ”-nr layers 4”. We select four layers because, in the next step, we apply the marching cubes algorithm as implemented in BrainVoyager to encapsulate the voxels with triangular tessellation. This means that the triangular mesh surfaces encapsulating the voxels corresponding to LayNii’s layers 1, 2, and 3 represent 25%, 50%, and 75% of the cortical depth, respectively.

Once LayNii outputs are generated, we convert the NIfTI files into BrainVoyager VMR format using bvbabel v0.1.0. Then, we reconstruct the triangular mesh surfaces using Brain-Voyager’s ”create mesh” function. Repeating this procedure for each layer results in three distinct triangular meshes. For example, in the right hemisphere of sub-01, the deep layer mesh contains 4,018,978 vertices and 8,037,952 triangles, the middle layer mesh has 4,239,784 vertices and 8,479,564 triangles, and the superficial layer mesh consists of 4,521,088 vertices and 9,042,172 triangles. As expected, the number of vertices increases from deep to superficial layers, reflecting the natural expansion of cortical surface area toward the outer cortex.

It is important to note that our varying vertex density meshes differ from conventional methods, where a single triangular mesh is first generated at the white matter boundary, and each vertex is projected outward along its normal vector by a fraction of the cortical thickness to form additional layers. This traditional method can result in uneven and insufficient coverage of underlying voxels due to the natural increase in surface area from deep to superficial layers and the fact that the human cortex has more gyri than sulci.

We deliberately opted for varying vertex density meshes to accurately represent geometric cortical layers. This choice is particularly crucial because the intracortical vessels we aim to highlight—mesoscopic ascending veins—appear as small circular structures on the reconstructed surfaces Adams et al. (2015) and Duvernoy et al. (1981). Therefore, it is essential to ensure that no underlying voxel is omitted due to suboptimal surface reconstruction, which could otherwise lead to missing critical vascular structures.

Once the layer surfaces were reconstructed, we first applied ”advanced mesh smoothing” in BrainVoyager using the following parameters: number of iterations = 150, smoothing force = 0.07, avoid shrinking = on. This step effectively smooths the initial blocky triangular tessellation while minimizing vertex displacement. Next, we used these meshes to sample voxel values using BrainVoyager’s ”Create SMP (Surface Map)” function with the ”sample volume data exactly at mesh vertices” option. This ensures that each vertex samples the value of a single voxel, rather than integrating values from neighboring voxels, preserving fine-scale vascular details. Following this, we applied ”mesh morphing” in BrainVoyager using the ”inflation mode”, running for 12,000 steps, to generate inflated surface renderings that enhance the visibility of sulcal intracortical vessels. Finally, we captured screenshots of the 3D renders after setting the surface shininess parameter to 1.0, ensuring minimal lighting effects for a clear and consistent visualization.

### 4.5 Data, Protocol, Code, and Software Availability

Our data, sequence parameters (PDFs), and part of the processed data used in will become available at: https://doi.org/10.5281/zenodo.14145584. The custom multi-shot 3D EPI sequence can be accessed via the Siemens C2P exchange platform. Processing and analysis scripts used in this study are available at: https://github.com/ofgulban/meso-MRI with 3DEPI. We used ITKSNAP v4.2.2 for the manual refinements on segmentations and cutouts. We used LayNii v2.7.0 using the LN2 LAYERS program for the “layerification” step. We used “real-time volume rendering” in our leptomeningeal and pial voxel value renders, and “triangular mesh reconstruction” in our intracortical mesh renders as implemented in BrainVoyager v24.0.

## Acknowledgements

O.F.G. and R.G. have financial interest tied to Brain Innovation. Scanning was supported by FPN (faculty of psychology and neuroscience) via the MBIC grant scheme. Scanning was performed at the facilities of Scannexus B.V. (Maastricht, Netherlands). O.F.G. acknowledges the passing of their grandfather, Hü seyin, during the preparation of this manuscript. His unwavering commitment to honesty and deep respect for science continue to serve as a lasting inspiration, shaping both this work and O.F.G.’s broader approach to scientific inquiry.

---

**Supplementary Figure 1:**
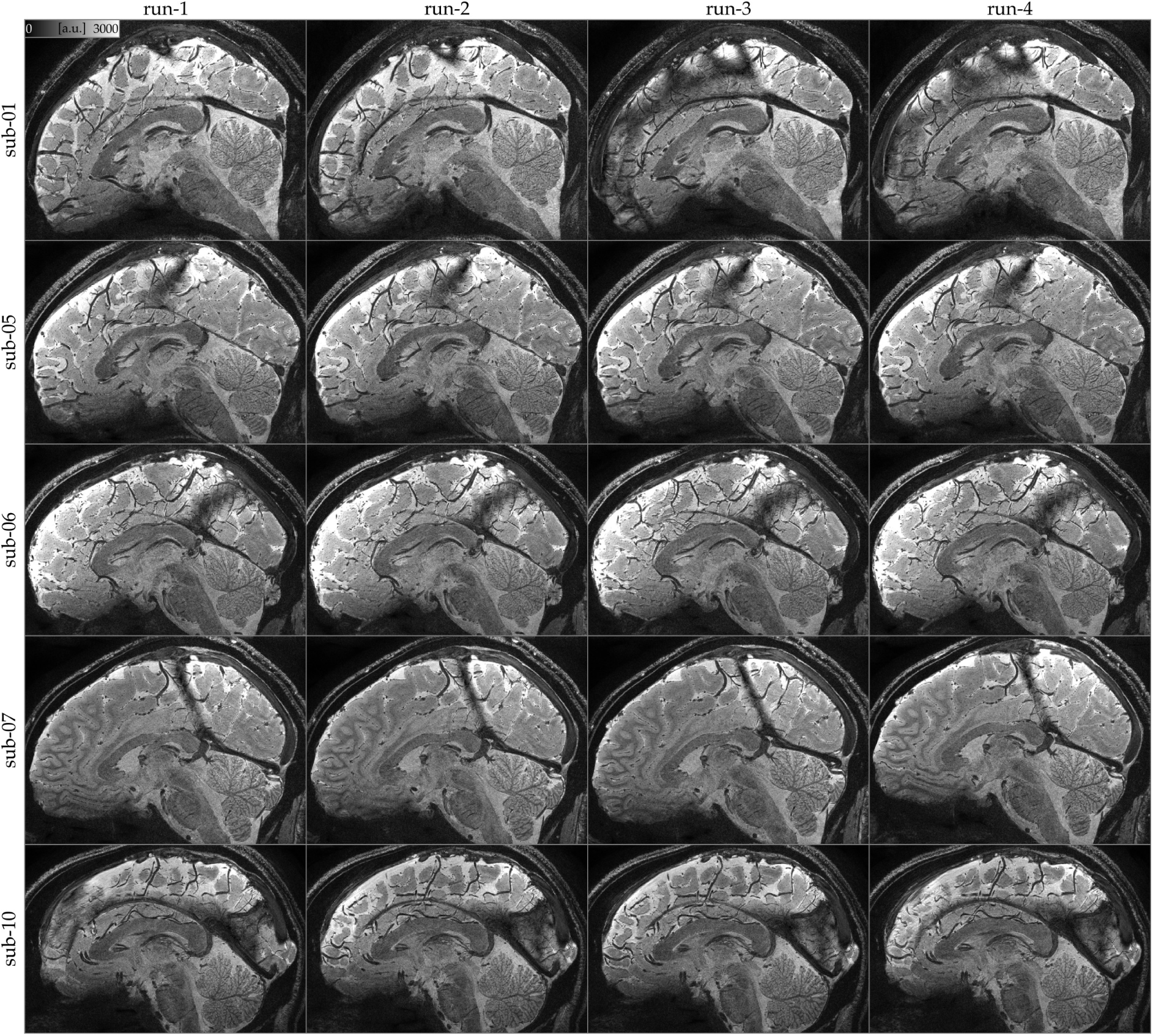
Imaging slab positioning and data quality. No data were discarded due to bulk head motion within acquisitions (see Gulban, et al. (2022), Supplementary Figure 3 for an example of within-run head motion). Notably, a substantial between-run head motion of nearly a centimeter (measured from the inferior frontal region) was observed in sub-07 run-04. However, this did not noticeably degrade image quality, particularly in resolving fine details of mesoscopic veins. While anecdotal, this finding highlights the effectiveness of our participant preselection strategy and the importance of explicit between-run instructions, such as encouraging participants to move if uncomfortable. Allowing controlled movement in between acquisitions increases the likelihood of obtaining high-quality images. Note that our images are acquired consecutively within a short period of time within 2024 and the gaps in subject numbers do not reflect any data removal.

**Supplementary Figure 2:**
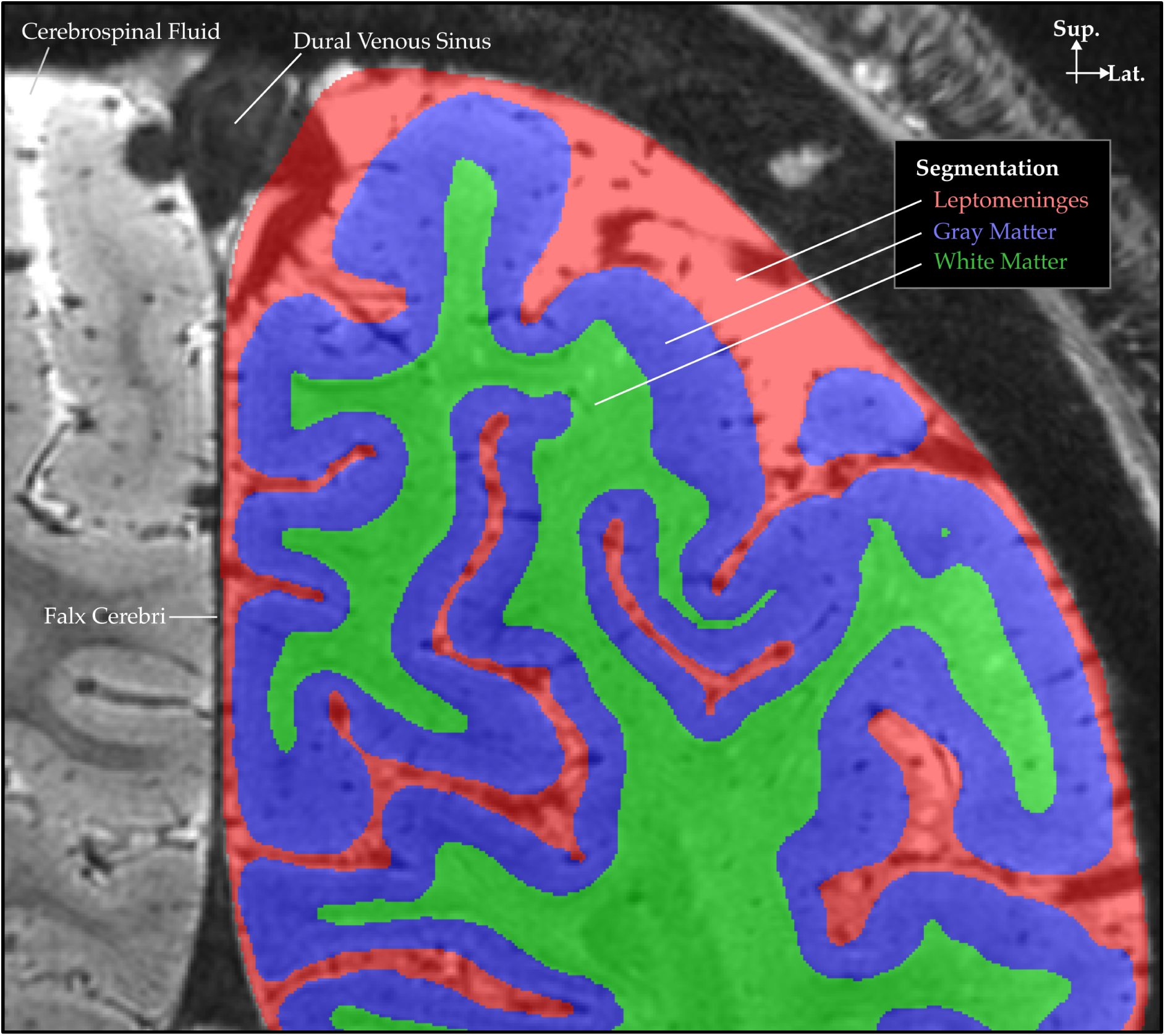
Segmented tissues overlaid on T_2_*-weighted images for sub-01’s left hemisphere. In our T_2_*-weighted images cerebrospinal fluid appears very bright, veins and large arteries appear very dark, cortical gray matter appears brighter than white matter. Manual edits on the initial segmentation were done using ITKSNAP v4.2.2 to improve the accuracy and precision of each tissue label. A conjunction mask of leptomeninges, gray matter, and white matter was applied to exclude other tissues before voxel value rendering in Brain-Voyager v24.0 (see Figure 5-7). Gray matter segmentation was used to compute geometric layers via the LN2 LAYERS program in LayNii v2.7.0, which were subsequently used to reconstruct triangular meshes in BrainVoyager (see Figures 8-9).

**Supplementary Figure 3:**
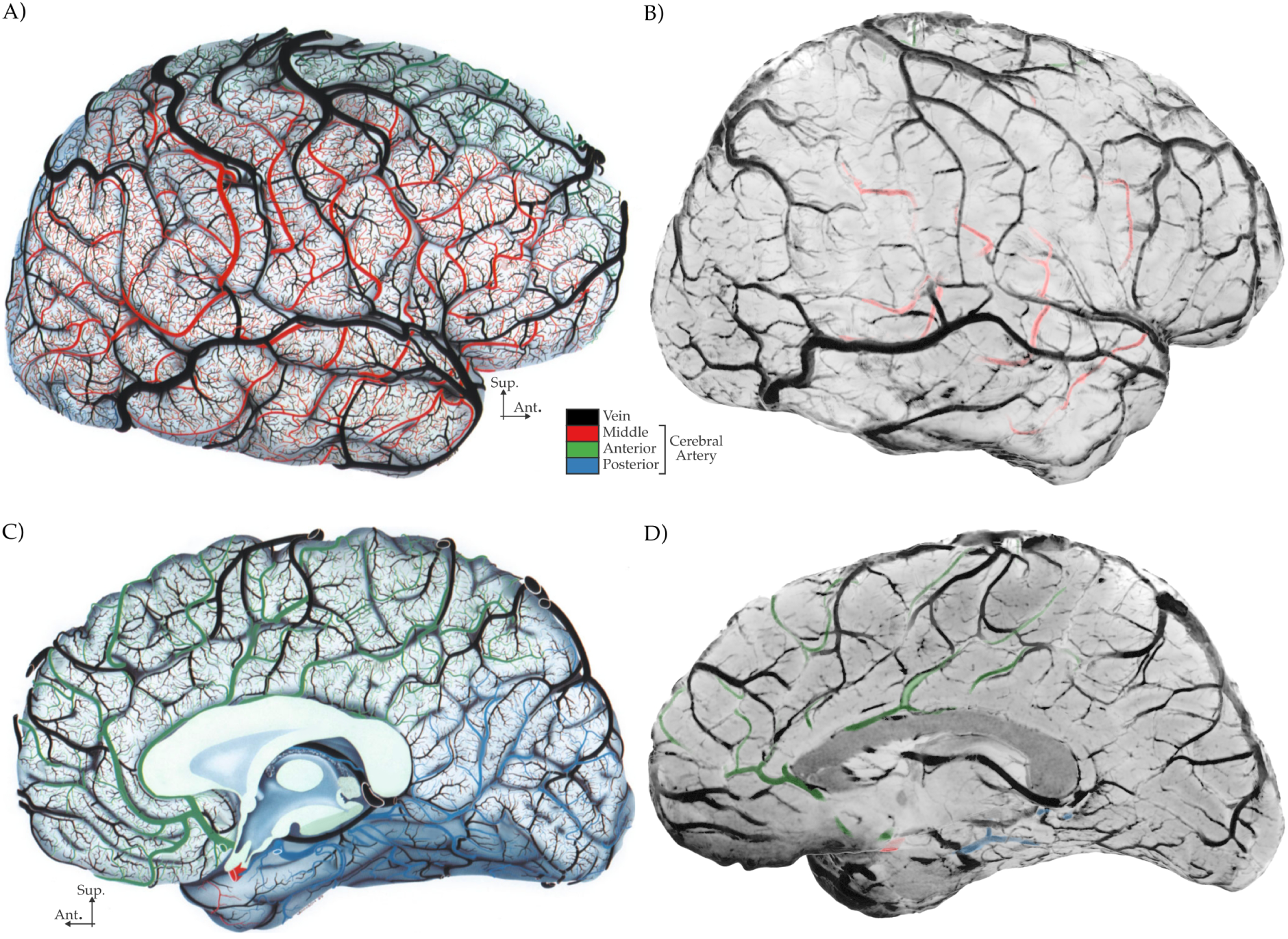
Comparison of cortical angioarchitecture visualized using post-mortem ink injection (Panels A and C, adapted from Duvernoy, et al. (1981)) and in vivo T_2_*-weighted MRI (Panels B and D). Note that smaller veins require a more zoomed in view to be visible on our in vivo images (see Figures 8-10). To be more comparable to Jean-Louis Vannson’s drawings, we inverted the intensity channel in our 1/T_2_*-weighted volume-rendered images, rendering veins dark and cortical gray matter bright, while color-coding large arteries. Panels A and B are adapted from Figures 233 and 234 in (Duvernoy & Vannson, 1999).

**Supplementary Figure 4:**
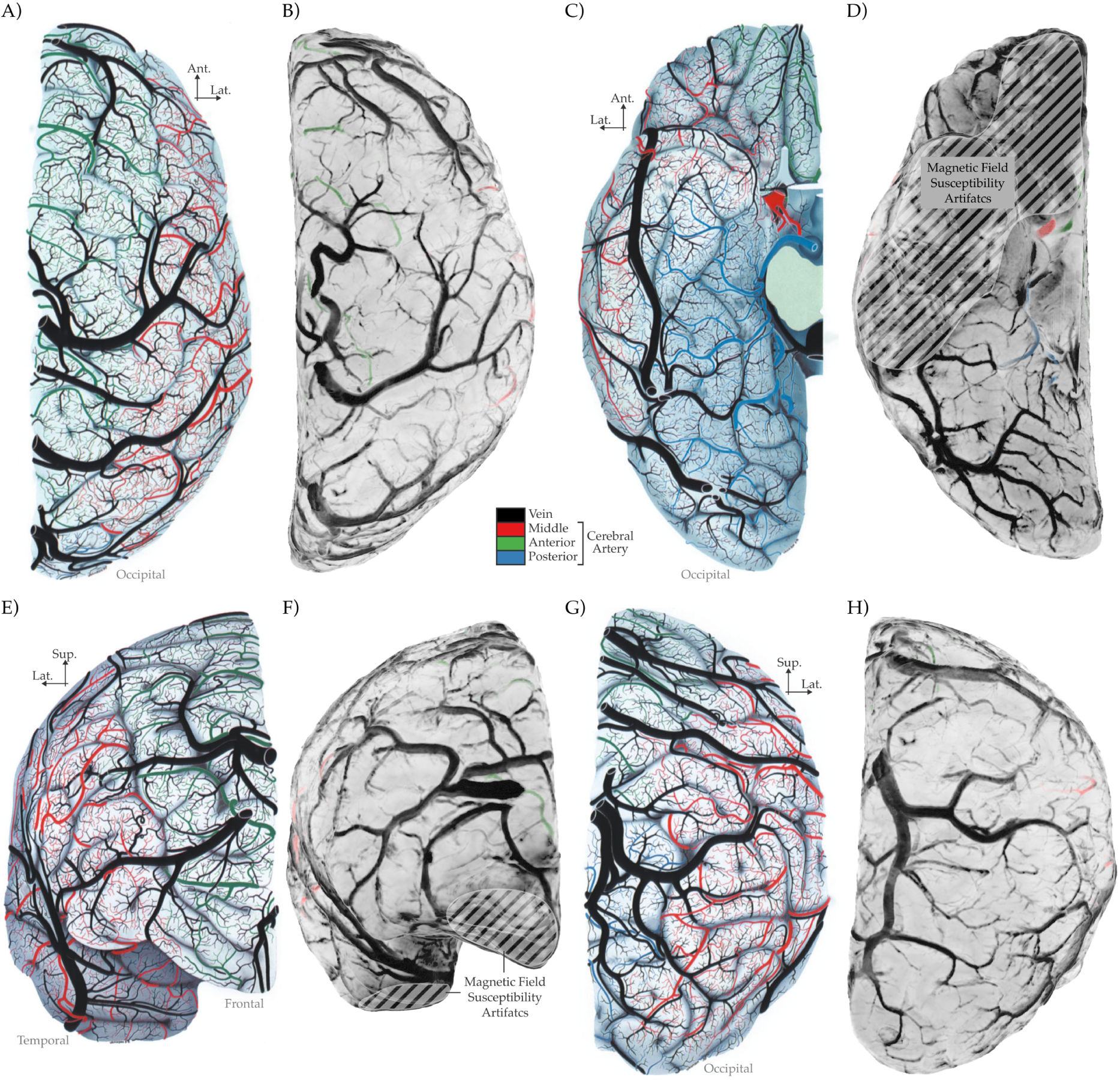
Comparison of comparing cortical angioarchitecture visualized using postmortem ink injection (Panels A, C, E, G, adapted from Duvernoy and Vannson (1999)) and in vivo T2*-weighted MRI (Panels B, D, F, H). Format same as Figure 5. In addition, we have marked the inferior brain regions affected by magnetic field susceptibility artifacts, where tissue segmentation becomes unreliable. Panels A, C, E, G are adapted from Figures 235-238 in (Duvernoy & Vannson, 1999).

**Supplementary Figure 5:**
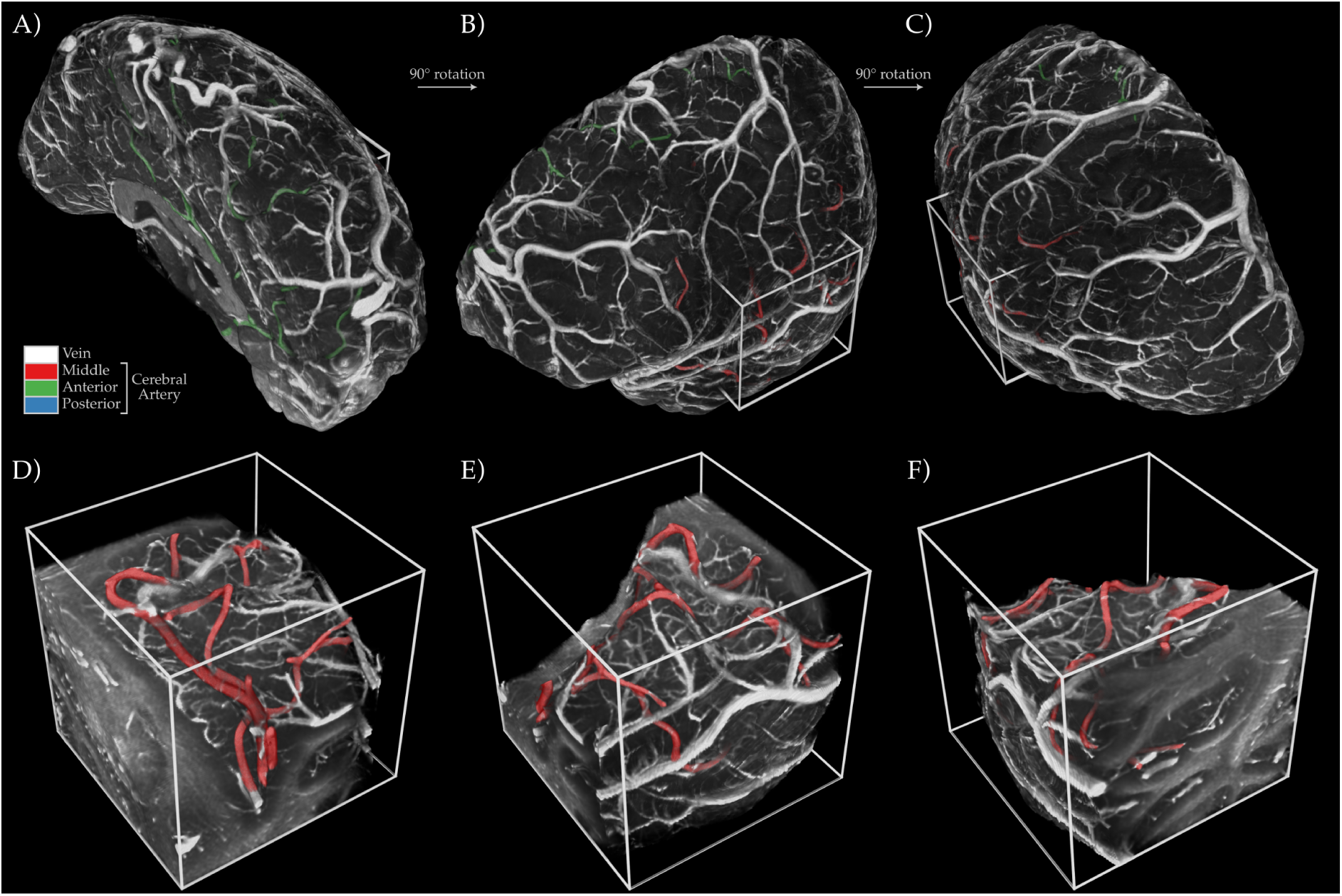
Different perspectives of the whole brain (A-B), and zoomed-in, cutouts (D-F) for temporal lobe adjacent vessels of Figure 7. Bright tubular structures are leptomeningeal and pial veins. The colored cerebral arteries are classified by tracing their branches towards the brainstem.

